# Inhibition of the Microglial Phagocytic Receptor MerTK Underlies ELA-induced Changes in Synapses and Behavior in Male Mice

**DOI:** 10.1101/2025.07.18.665641

**Authors:** Madison M. Garvin, Urjoshi Kar, Cassandra L. Kooiker, Jessica L. Bolton

**Affiliations:** Neuroscience Institute, Georgia State University, Atlanta, GA, USA; Departments of Pediatrics and Anatomy/Neurobiology, University of California, Irvine, Irvine, CA, USA

**Keywords:** microglia, MerTK, synaptic pruning, PVN, CRH+ neurons, stress, threat response, early-life adversity

## Abstract

**Background:** Early-life adversity (ELA) is a significant risk factor for emotional disorders like depression, likely by provoking changes in stress-related circuit development. We have previously shown that ELA increases the number of excitatory synapses onto corticotropin-releasing hormone (CRH)-expressing neurons in the paraventricular nucleus (PVN) by decreasing microglial synapse engulfment. Here, we hypothesize that ELA induces microglial dysfunction via inhibition of the microglial phagocytic receptor, MerTK, thus resulting in the observed changes in synapses and stress-related behavior.

**Methods:** To determine whether deleting MerTK in microglia phenocopies the effects of ELA, microglia-specific (m)MerTK-KO (CX3CR1-Cre^+^::MerTK^fl/fl^) mice were crossed with ‘wild-type’ (CX3CR1-Cre^-^::MerTK^fl/fl^) mice and their litters were reared in either a control or ELA (induced by limited bedding and nesting paradigm) environment, from postnatal days (P)2-10. Excitatory synapses in the PVN were assessed at P10, and adult offspring were tested in a behavioral battery to measure threat-response (known to be dependent on PVN-CRH+neurons) and anxiety-like behavior, followed by acute restraint stress to measure the neuroendocrine stress response.

**Results:** Following ELA at P10, excitatory, but not inhibitory, synapses in the PVN were increased in males, which was mimicked by mMerTK-KO in control males, but caused no further increase in ELA males. However, females already had higher numbers of excitatory synapses at baseline, and showed no further increase with ELA or mMerTK-KO. Remarkably, the pattern of threat-response behavior in males closely matched the excitatory synapses, with mMerTK-KO control males escaping more from the simulated predator threat in the looming-shadow threat task, similar to ELA males. Again, females did not show any significant changes due to ELA or mMerTK-KO in the threat-response, although they did show ELA-induced changes in anxiety-like behavior. ELA provoked a greater corticosterone response to acute stress in males, but not females, although females were again higher at baseline.

**Conclusions:** Our results demonstrate that ELA provokes increased excitatory synapses in the PVN, leading to an increased active response to threat in the looming-shadow test in males only. Deleting MerTK specifically from microglia recapitulates both the synaptic and behavioral effects in control males, but does not have an effect in ELA males or control females, suggesting that the MerTK pathway is already inhibited by ELA in males and less active in females at baseline. Our work is the first to elucidate the mechanisms underlying the male-biased microglial dysfunction caused by ELA, with promise for the development of better preventative and therapeutic strategies for at-risk children.

## 1. Introduction

Early-life adversity (ELA) is a robust risk factor for neuropsychiatric disorders like depression, which are often characterized by abnormal stress responses and a dysregulated hypothalamic-pituitary-adrenal (HPA) axis (Bick & Nelson, 2016; Danese & McEwen, 2012; Heim & Binder, 2012; McLaughlin et al., 2019; Stout & Nemeroff, 1994; Watson & Mackin, 2006). ELA can profoundly impact how an individual responds to future stressors by modulating the maturation of stress-responsive brain circuits, and their fundamental building blocks, synapses (Bolton et al., 2018, 2022). However, the mechanisms by which ELA modulates synapse number and circuit development remain poorly understood.

It is now well-established that glial cells, and particularly microglia, the brain’s resident immune cells and primary phagocytes, prune synapses during brain development, which is critical for the normal maturation of neurons and their circuits (Paolicelli et al., 2011; Schafer et al., 2012; Weinhard et al., 2018). Synaptic refinement via microglial engulfment is especially susceptible to the effects of ELA, which we have previously investigated using a limited bedding and nesting model during the first week of life (Bolton et al., 2022). For example, we have found that ELA in the first week of life increases the number of excitatory synapses onto corticotropin-releasing hormone (CRH)-expressing neurons in the paraventricular nucleus of the hypothalamus (PVN) by inhibiting microglial synapse engulfment (Bolton et al., 2022). Our recent work has shown that ELA provokes an aberrant stress response, as measured by the neuroendocrine response to an acute stressor and the escape response to the looming-shadow threat task (Bolton et al., 2022), both of which require PVN-CRH+ neurons (Daviu, Füzesi, Rosenegger, Rasiah, et al., 2020).

Intriguingly, we also found that all of the above-described effects of ELA are greater in males than females. In humans too, neuroanatomical changes after ELA, such as smaller hippocampal volumes associated with depression, are often more profound in men than women (Colle et al., 2017; Teicher et al., 2004), and males across species are more vulnerable to ELA during the perinatal sensitive period (McCarthy, 2016). However, the mechanistic basis of the sex-specific effects of ELA are unknown, and here, we aim to address this important question.

Microglia and astrocytes prune synapses in response to several cellular and molecular triggers (Faust et al., 2021), including activation of the phagocytic receptor Mer tyrosine kinase (MerTK), which recognizes externalized phosphatidylserine on the surface of whole cells and parts of cells (like synapses), that need to be cleared (Abiega et al., 2016; Diaz-Aparicio et al., 2020; Fourgeaud et al., 2016). This pathway has recently been shown to interact with the complement system, another important “eat-me” signal that tags synapses for removal by microglia (Scott-Hewitt et al., 2020). MerTK has been mainly investigated thus far in astrocytic synaptic pruning mechanisms (Chung et al., 2013), despite its expression by both microglia and astrocytes in the brain (Zhang et al., 2014). The relative contributions of microglia vs. astrocytes to synaptic pruning are still debated in the field, and the controversy may be complicated by differences based on brain region and sex. In our prior work, we have observed MerTK expression predominantly in microglia in the immature PVN, and we have identified MerTK as a potential molecular pathway by which ELA impairs microglial synaptic pruning of PVN-CRH+ neurons in males (Bolton et al., 2022). We reported that ELA decreases microglial MerTK expression in males in the P8 PVN (Bolton et al., 2022), but in females, we now find that MerTK is already lower at baseline and not altered by ELA (Figure 1A), in agreement with their diminished synaptic pruning (Garvin & Bolton, 2022). Furthermore, we found that pharmacological inhibition of MerTK *in vitro* increases the number of excitatory synapses onto CRH+ neurons in control male slice cultures to the level of ELA, but fails to further augment synapse number in ELA male cultures (Figure 1B) (Bolton et al., 2022) or cause any significant changes in control female cultures (Figure 1C). Together, these data suggest that the microglial MerTK pathway is already inhibited by ELA (i.e., an occlusion effect) in males, but not significantly altered by ELA in females, perhaps due to a lower baseline activity.

**Figure 1.**
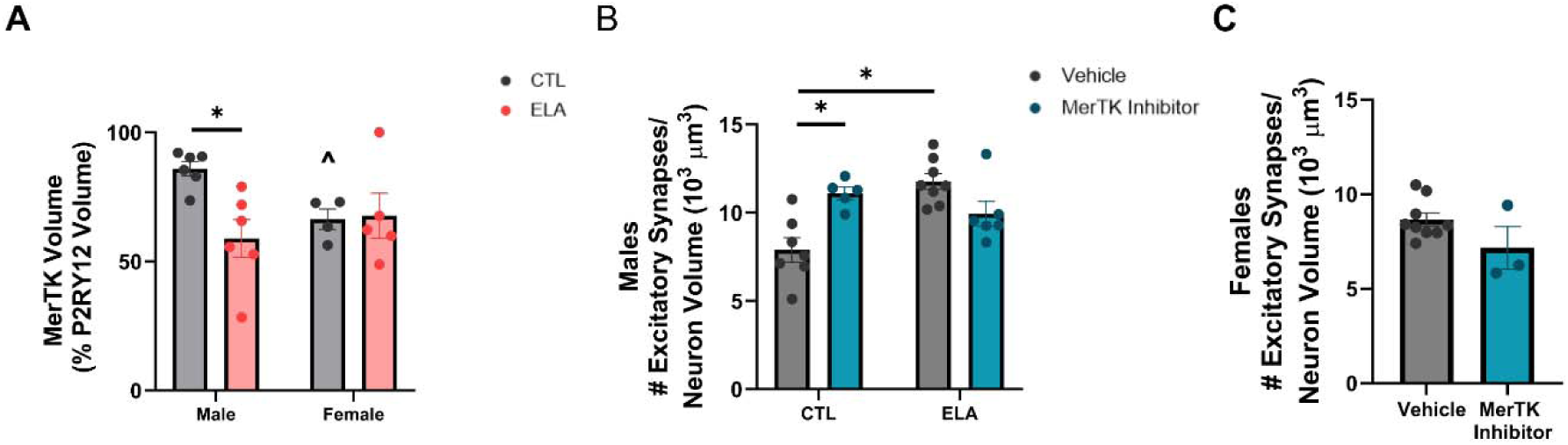
Evidence identifying MerTK as a potential mechanism underlying the ELA-induced increase in excitatory synapses. (A) ELA reduces MerTK volume per volume of P2RY12+ microglia in the PVN of P8 male mice (Adapted from (Bolton et al., 2022), whereas females already have lower MerTK levels at baseline and no further decrease due to ELA (significant Sex x ELA interaction, *F*_1, 17_ = 4.931, *p* = 0.0403; 2-way ANOVA; *p*<0.05; post hoc Šídák’s test). (B) Pharmacological application of MerTK inhibitor in PVN organotypic slice cultures obtained from control males increases the number of excitatory synapses to the level of ELA+vehicle males, but does not further increase synapse number in ELA conditions (significant Treatment x ELA interaction, *F*_1, 22_ = 17.89, *p*= 0.0003; 2-way ANOVA; p<0.05; post hoc Šídák’s test (Adapted from (Bolton et al., 2022)). (C) However, application of MerTK inhibitor does not significantly alter the number of excitatory synapses in PVN slice cultures from control females (unpaired *t*-test, *p*>0.1), similar to the lack of effect of ELA we have previously reported (Bolton et al., 2022). Data are means ± SEMs; \**p*<0.05, ^*p* = 0.1, post hoc Šídák’s test, following up a Sex x ELA or ELA x Treatment interaction.

Here, we aim to test the hypothesis that MerTK is the molecular pathway by which ELA impairs microglial synaptic pruning in males, but not females. There are important limitations for our prior published data that we aim to address here: first, it was *in vitro,* and we now know microglia are fundamentally altered in cultures due to their exquisite sensitivity to the microenvironment (Gosselin et al., 2017). Secondly, our pharmacological treatments could have impacted both microglial and astrocytic MerTK. Thus, here we have conducted all *in vivo,* cell type-specific experiments in order to best test our hypothesis.

## 2. Material and Methods

### 2.1 Animals

All mice were housed in quiet, uncrowded conditions on a 12-hr light/dark cycle with *ad libitum* access to food and water. The transgenic mouse lines employed in this study were on a C57BL/6 background and include CX3CR1-BAC-Cre^+^ (GENSAT; MMRRC stock #036395-UCD; obtained from Dr. Staci Bilbo, Duke University), and MerTK^fl/fl^ (obtained from Dr. Carla Rothlin, Yale University). CX3CR1-Cre^+/-^::MerTK^fl/fl^ mice were bred with CX3CR1-Cre^-/-^::MerTK^fl/fl^ mice to obtain experimental animals (Figure 2). CRH-Cre^+/-^ mice (Jax #: 012704; Jackson Laboratory, Bar Harbor, ME) were crossed with tdTomato^+/+^ (stop-floxed; Ai14; Jax #: 007914) to generate CRH-reporter mice for Figure 1 and Supplemental Figure 1. Both male and female offspring were used throughout the entire study. All experiments were performed in accordance with National Institutes of Health (NIH) guidelines and were approved by the Georgia State University Animal Care and Use Committee.

**Figure 2.**
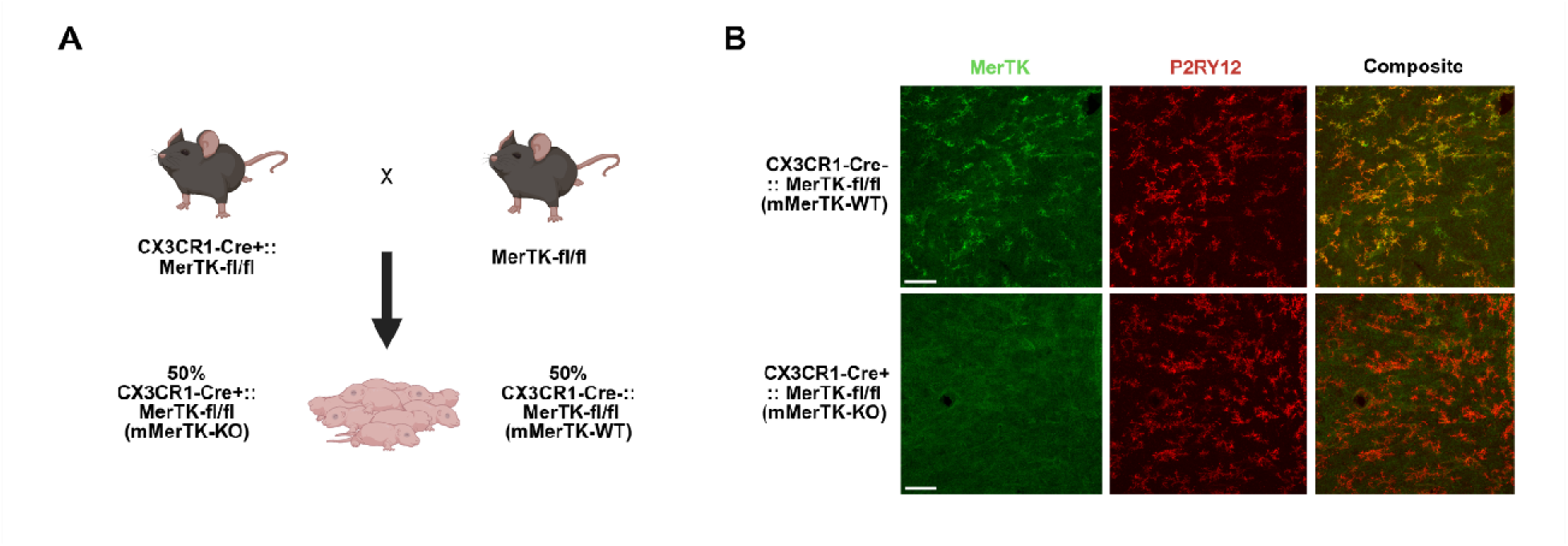
Validation of microglia-specific MerTK deletion (mMerTK-KO) *in vivo*. (A) Breeding scheme for *in vivo* studies: CX3CR1-Cre^+^::MerTK^fl/fl^ mice were crossed with MerTK^fl/fl^ mice to generate ∼50% of pups in each litter with MerTK specifically knocked out from microglia, and ∼50% with ‘wild-type’ levels of microglial MerTK. The diagram was created with BioRender.com. (B) Representative images of MerTK expression colocalizing with P2RY12 (microglia) in the PVN of mMerTK-WT (top) vs mMerTK-KO (bottom) mice, validating the microglia-specific MerTK deletion. Scale bar= 50 μm.

### 2.2 Limited bedding and nesting (LBN) model of early-life adversity (ELA)

Experimental dams were checked daily for copulatory plugs and then single-housed in our quiet, temperature-controlled maternal care room on embryonic day (E)17. On the morning of postnatal day (P)2, litters were culled to 8 pups and dams and their litters were randomly assigned to either a control or early-life adversity (ELA) group. Dams in the control group were transferred to a standard cage with ∼220 grams of corn cob bedding and one full cotton-square nestlet. Dams in the ELA group were transferred to a cage fitted with a mesh platform (25 x 19 cm, cat # 4700313244; McNichols Co., Atlanta, GA) sitting approximately 2.5 cm above the floor. Bedding was reduced to ∼110 grams and half of a cotton square nestlet was provided (Mroue-Ruiz et al., 2024). The cages for both groups remained undisturbed from P2 until P10 when the dams and pups were transferred back to standard cages in the colony room for growing up to adulthood for behavioral testing or P10 pups were euthanized to assess excitatory synapses.

### 2.3 Method Details

#### 2.3.1 Immunohistochemistry (IHC) for synaptic markers

P10 mice of both sexes were euthanized with Euthasol and perfused transcardially with 1x phosphate-buffered saline (PBS; pH = 7.4) followed by 4% paraformaldehyde in 0.1 sodium phosphate buffer (pH = 7.4). Brains were post-fixed for 4 hours and then transferred to 25% sucrose overnight for cryoprotection. Tissue was frozen and sectioned coronally into 25-µm-thick slices using a Leica CM1860 cryostat. For immunohistochemistry, floating brain sections were washed 3 times in PBS-T (PBS containing 0.3% Triton X-100, pH = 7.4) and then incubated in 0.9% H_2_0_2_ in PBS-T for 20 min. at room temperature (RT). To prevent non-specific binding, tissue was then incubated in a blocking solution containing 5% normal donkey serum (NDS) (cat# 017-000-121, Jackson ImmunoResearch, West Grove, PA, USA) for 1 hr. For excitatory synapse labeling, sections were then incubated at 4°C overnight with guinea pig anti-vGlut2 antiserum (1:12,000, cat# AB2251-1, Millipore Sigma, Temecula, CA, USA) and rabbit anti-PSD95 antiserum (1:1,000; cat#51-6900, Invitrogen/ ThermoFisher, Waltham, MA) in a PBS-T solution containing 2% NDS. The following day, sections were washed 3 times with PBS-T and incubated in the secondary antibodies, donkey anti-guinea pig IgG-594 (1:1,000, cat# 706-585-148, Jackson ImmunoResearch) and donkey anti-rabbit IgG-488 (1:1,000; A-21206, Invitrogen/ThermoFisher) in 2% NDS for 3 hr. at RT. Sections washed a final 3 times in PBS-T and were then stained with DAPI, mounted on gelatin-coated slides and coverslipped with Fluoromount-G (cat# 00-495802, Invitrogen/ThermoFisher). For inhibitory synapse labeling, sections were incubated with the primary antibodies, rabbit anti-vGAT antiserum (1:10,000, cat# 131 013, Synaptic Systems, Göttingen, Germany) and guinea pig anti-Gephyrin antiserum (1:500; cat# 147-318, Synaptic Systems). The corresponding secondary antibodies were donkey anti-rabbit IgG-488 (1:1,000; A-21206, Invitrogen/ ThermoFisher) and donkey anti-guinea pig IgG-594 (1:250; cat# 706-585-148, Jackson ImmunoResearch).

##### 2.3.1.1 Confocal imaging and analysis

Images of the mediodorsal parvocellular (mpd) paraventricular hypothalamic nucleus (PVN) were collected using a Zeiss LSM-780 confocal microscope (Zeiss, Dublin, CA, USA) with a 63x oil objective as previously (Bolton et al., 2022). 24 z-planes of 134.8 x 134.8 μm were taken at 0.5-µm intervals, and the image frame was digitized at 12-bit using a 1024 x 1024-pixel frame size. We were unable to use CRH-tdTomato+ mice here as previously (Bolton et al., 2022) to identify CRH+ neurons specifically due to already using a conflicting Cre (CX3CR1), and CRH IHC is notoriously inadequate for labeling whole neurons (Yan et al., 1998). Thus, an ROI was manually drawn around the neurons in the mpd PVN based on the densest area of DAPI staining, which we find to reliably overlap with the dense hub of PVN-CRH+ neurons (Supplemental Figure 1). Excitatory or inhibitory synapses were assessed as colocalized puncta of either vGlut2 + PSD95 or vGAT + Gephyrin, respectively, within the ROI volume, using Imaris’ (Bitplane, Zurich, Switzerland) colocalization function (threshold = 0.5).

#### 2.3.2 IHC for MerTK

Similar to the IHC procedure described above, coronal sections of adult tissue were washed 3 times in PBS-T and then incubated in 0.9% H202 in PBS-T for 20 minutes at RT. Following several washes, free-floating sections were then permeabilized in 0.5% Triton X-100 in PBS for 10 minutes. After 3 more washes in PBS-T, sections were blocked in 5% NDS for 1 hour, followed by incubation in rat anti-MerTK (1:1,000, cat# 14-5751-82, eBioscience) and rabbit anti-P2RY12 (1:2,000, cat# AS-55043A, AnaSpec,) in 2% NDS in PBS-T at 4°C overnight. The following morning, sections were washed 3 times in PBS-T and then incubated in donkey anti-rat IgG-488 (1:500, A-21202, Invitrogen Life Technologies) and donkey anti-rabbit IgG-647 (1:1,000, cat# 711-605-152, Jackson ImmunoResearch) in 2% NDS for 3 hours at RT. Sections were then washed a final 3 times in PBS-T, stained with DAPI (4’,6-diamidino-2-phenylindole), mounted on gelatin-coated slides and coverslipped with Fluoromount-G (cat# 00-495802, Invitrogen/ ThermoFisher).

##### 2.3.2.1 Confocal imaging and analysis

Images of the mediodorsal parvocellular (mpd) paraventricular hypothalamic nucleus (PVN) were collected using a Zeiss LSM-780 confocal microscope with a 20x objective. 12 z-plane images of 320.09 x 320.09 μm were taken at 1-µm intervals, and the image frame was digitized at 12-bit using a 1024 x 1024-pixel frame size. ROIs were manually drawn around the PVN as a whole (Supplemental Figure 1), and microglial MerTK volume was analyzed via Imaris 3D-reconstruction software and divided by P2RY12 volume.

#### 2.3.3 MerTK inhibitor treatment of organotypic PVN slice cultures

Following the rapid decapitation of P6-7 male CRH-Cre+/-:tdTomato+/- mouse pups, the brains were dissected from the skull, and hypothalamic blocks were removed and cut into 350-mm coronal sections on a McIlwain tissue chopper. Approximately 6 sections were then collected posterior to the anterior commissure in order to create the organotypic PVN cultures. These slices were explanted onto Millicell cell culture inserts (pore size 0.4 mm, diameter 30 mm, Merck Millipore Inc., cat# PICMORG50). Membrane inserts were placed into a six-well plate with 1 mL of culture medium. Culture medium consisted of 52% modified Eagle’s medium [MEM; cat# 11700, Invitrogen], 25% Hanks Balanced Salt Solution [HBSS; cat# 24020, Invitrogen], 20% Heat Inactivated Horse Serum (added post-filtration) supplemented with 3 mM L-Glutamine (Gibco, cat# 25030081), 25 mM D-Glucose (Sigma, cat# G7528), 1.9 mM NaHCO3 (ThermoFisher, cat# 25080094), 12.5 mM HEPES (Gibco, cat# 15630080), 0.6 mM L-Ascorbic acid (Sigma-Aldrich, cat# A4403), 1 mg/mL Insulin (Sigma-Aldrich, cat# I0516) and 25 mg/mL containing penicillin-streptomycin (Gibco, cat# 15140122). Slices were cultured at 37°C in 5% CO2 enriched air for 48 hr., at which point the medium was refreshed. Medium without antibiotics was then used to refresh the media after an additional 48 hours. 6 days after initial culture, each membrane was washed twice with 2 mL of antibiotic-free, serum-free medium (97% MEM supplemented with 3 mM L-Glutamine, 10 mM D-Glucose,1.9 mM NaHCO3, 12.5 mM HEPES, 0.6 mM L-Ascorbic acid, 1 mg/mL Insulin) and treatment with 20 nM of a small-molecule MerTK inhibitor as previously described (UNC2025, Selleck Chemicals, cat# S7576) (Zhang et al., 2014a) or sterile culture-grade water as vehicle was commenced. UNC2025 has IC50 = 2.7 nmol/L and 40-fold greater selectivity for MerTK over Axl and Tyro3 (McDaniel et al., 2018). UNC2025 also acts on FLT3, but this molecule is mostly expressed in hematopoietic stem cells with no expression in the neonatal mouse brain (Ito et al., 1993). Media was then refreshed 12 hr. later with serum-free media containing either vehicle or MerTK inhibitor. After an additional 4 hr., cultures were fixed in 4% paraformaldehyde in 0.1 M phosphate-buffer (PB) on ice for 30 min. rinsed in PB, and cryoprotected in a 25% sucrose solution for 4–6 hr.

#### 2.3.4 Organotypic PVN slice cultures IHC and imaging

Cultures that were confirmed to contain a PVN were individually frozen on dry ice while still attached to the membrane. Slice cultures were sectioned to 14-mm thickness using a Leica CM1900 cryostat and mounted immediately onto gelatin-coated slides. Slides were stored at 20°C until the IHC procedure. After thawing, slides were rinsed in PBS-T, followed by treatment in 0.3% H_2_O_2_ in 0.01 M PBS for 20 min. Sections were blocked in PBS-T containing 5% NDS and then incubated overnight at 4°C with rabbit anti-PSD95 (1:1000, Invitrogen/ThermoFisher) and guinea pig anti-vGlut2 (1:10,000, Millipore Sigma) in PBS-T containing 2% NDS. Sections were washed in PBS-T and then incubated in donkey anti-guinea pig IgG conjugated to Alexa-Fluor 647 (1:500, Jackson ImmunoResearch) and donkey anti-rabbit pig IgG conjugated to Alexa-Fluor 488 (1:500, ThermoFisher) in PBS-T containing 2% NDS for 3 hr. at room temperature. Sections were washed with PBS-T and coverslipped with Vectashield mounting medium with DAPI (Vector Laboratories). Confocal images of the mpd PVN were collected with an LSM-510 confocal microscope (Zeiss) with an Apochromat 363 oil objective. 11 z-stack images of 142.86 x 142.86 mm were taken at 1-mm intervals. Image frame was digitized at 12-bit using a 1024 3 1024-pixel frame size (Figure S3E). CRH neuronal volume was automatically calculated using Imaris’ 3D reconstruction function. Excitatory synapses onto CRH+ neurons were identified as colocalized puncta of vGlut2+PSD95 within the CRH-tdTomato+ volume using Imaris’ colocalization function (threshold = 1.0).

#### 2.3.5 Behavioral assays

Adult (>P60) mice were transferred to a 12-hour reverse-light cycle room with *ad libitum* access to food and water at least one week before initiation of behavioral testing. All tasks were completed in a quiet room illuminated with dim, red lighting in which subjects were acclimatized for 1 hr. in their home cage before any behavioral manipulation. Each apparatus was cleaned with 10% ethanol between subjects to avoid residual odor cues. Testing and analyses were conducted by an experimenter who was blinded to treatment group. All behavioral data were analyzed using the means for each litter, in order to avoid litter effects.

##### 2.3.5.1 Looming-Shadow Threat (LST) Task

Before testing, mice were habituated for 15 min. to an arena (43 cm x 18 cm x 16 cm) containing a plastic, matte black shelter in one corner. The stimulus, consisting of a black disk that was stationary for 3 s, expanded for 2 s and then stationary for another 3 s, was then presented to the subject 5 times with a separation period of at least 1 minute and 15 seconds between each presentation (Daviu, Füzesi, Rosenegger, Peringod, et al., 2020). Responses to the stimulus were manually scored as no response, freezing, or escaping, which was then verified later with a video recording. The scoring window was 14 seconds, consisting of the 8 s stimulus plus an additional 6 seconds to account for delayed escape behavior to make it into the shelter. The criteria for scoring behavior were defined as follows: 1) a flight response resulting in the subject entering or reaching the edge of the shelter within 14s of the onset of the stimulus was classified as escape; 2) the total suspension of movement aside from breathing was classified as freezing; 3) all other behaviors or lack thereof were classified as no response. If both freezing and escape behavior occurred in the same trial, escape behavior was scored as it is the more active of the two behaviors. The latency to respond to the stimulus was also measured, with the latency for no response being recorded as 14 seconds.

##### 2.3.5.2 Open-Field Test (OFT)

Following a one-week resting period after LST, a 10-minute open-field test was conducted in an arena (43LcmL×L43Lcm x 30 cm, cat# ENV-515S-A, Med Associates Inc, Fairfax, VT, USA) encased in a white opaque, sound-proof box with double doors (cat# ENV-017, Med Associates Inc). Using Med Associates tracking software (Med Associates Inc), the arena was divided into 225 equivalent squares with the 144 outermost squares classified as the “surround” zone and the 81 innermost squares classified as the “center” zone. The percentage of time spent in the center zone was used as a measure of anxiety-like behavior, and the total distance traveled was used to indicate locomotor activity (Simon et al., 1994).

##### 2.3.5.3 Elevated-Plus Maze (EPM)

Following a rest period of at least 3 days after the OFT, a 5-minute elevated-plus maze test was conducted in an apparatus consisting of two closed arms (35 cm × 5 cm × 15 cm; black opaque walls) and two open arms (35 cm × 5 cm × 0.5 cm) crossing perpendicularly and raised 50 cm off the floor. Any-Maze tracking software (Stoelting Co., Wood Dale, IL, USA) was used to measure the total distance travelled to indicate locomotor activity and the percentage of time spent in the open arms to assess anxiety-like behavior. The percentage of time spent in the open arms was calculated as the time spent in the open arms divided by the time spend in the open and closed arms, thus excluding the time spent in the intersecting center (Seillier & Giuffrida, 2017).

#### 2.3.6 Acute stress testing in adulthood

Adult mice were placed in a restraining device created from a 50-mL conical tube (cat# 339652, ThermoFisher) for 90 min. Blood samples (∼100 µl) were collected from the facial vein using a 4mm lancet (cat# NC9922361, Medipoint/Fisher Scientific) at baseline, 15 min., 30 min., and 60 min. after the initiation of the stress period. Following a 30 min. period at room temperature to allow for blood clotting, samples were centrifuged at 3000 x *g* for 15 min., and serum was collected and then stored at -20°C until later analysis. Serum was analyzed for CORT using an enzyme-linked immunosorbent assay kit according to the manufacturers’ instructions (cat# 501320, Cayman Chemical).

### 2.4 Quantification and Statistical Analysis

Statistical analyses were all performed using GraphPad Prism 10.0 software (GraphPad, San Diego, CA, USA). Three-way ANOVAs (Sex x ELA x Genotype) were used to analyze all data, except in Figure 1, where 2-way ANOVAs (Fig. 1A-B) and an unpaired *t-*test were used (Fig. 1C). Following a significant or trending (p≤0.1) interaction with Sex, 2-way ANOVAs within males and females were used to analyze the effects of ELA and genotype. Any interactions of ELA x Genotype were followed by Šídák’s post hoc tests. For the sake of consistency, all data were separated by sex in the graphs due to the observation of multiple sex differences. Data are presented as mean ± SEM and Grubbs’ test was used to remove statistical outliers (alpha=0.05) from the data. All experimental analyses were conducted in a blinded manner, with the experimenter having no knowledge of experimental group.

## 3. Results

### 3.1 Microglia-specific deletion of MerTK augments the number of excitatory synapses in the PVN of control males to the level of ELA males

Our previous findings have shown that inhibiting MerTK *in vitro* increased the number of excitatory synapses onto mediodorsal parvocellular (mpd) PVN neurons in organotypic slice cultures from males, mimicking the effect of ELA (Figure 1B) (Bolton et al., 2022). However, this effect was not seen in slice cultures from females (Figure 1C), which may be due to their already lower level of MerTK (Figure 1A) and higher number of excitatory synapses that does not change with ELA (Bolton et al., 2022). In order to more conclusively define the role of microglial MerTK in the observed ELA-induced alterations to synapse development *in vivo*, we generated a conditional knockout (KO) mouse line in which MerTK is specifically deleted from microglia (Figure 2A-B). Offspring were then reared in either control or ELA conditions between postnatal days (P)2 and P10 as previously described (Mroue-Ruiz et al., 2024), at which time the number of excitatory and inhibitory synapses in the mpd PVN were quantified (Figure 3A). As seen previously, ELA increased the number of excitatory synapses onto mpd PVN neurons in male mice (significant Genotype x ELA interaction, *F*_1, 30_ = 6.059, *p*=0.02 ; 2-way ANOVA; *p* < 0.05; post hoc Šídák’s test; Figure 3C). This effect was recapitulated by the microglia-specific MerTK KO (mMerTK-KO) in control males (*p*<0.05; post hoc Šídák’s test; Figure 3C). Notably, there was no further increase in synapses due to mMerTK-KO in ELA males, indicating that this pathway may already be inhibited by ELA (Figure 3B, C). As expected based on Figure 1, the mMerTK-KO did not alter excitatory synapse counts in female mice, although their numbers are already higher at baseline (significant main effect of Sex, *F*_1, 63_ = 32.13, *p*<0.0001; trend for Genotype x ELA x Sex interaction, *F*_1, 63_ = 2.994, *p*=0.09; 3-way ANOVA; Figure 3D). Furthermore, there were no significant effects of ELA or mMerTK-KO on the number of inhibitory synapses in the mpd PVN of male or female mice, although females had a trend for an ELA-induced decrease in inhibitory synapses (trend for ELA x Sex interaction, *F*_1, 62_ = 2.913, *p*=0.09; 3-way ANOVA; trend for main effect of ELA, *F*_1, 29_ = 4.106, *p*=0.05; 2-way ANOVA within females; Figure 2E-F).

**Figure 3.**
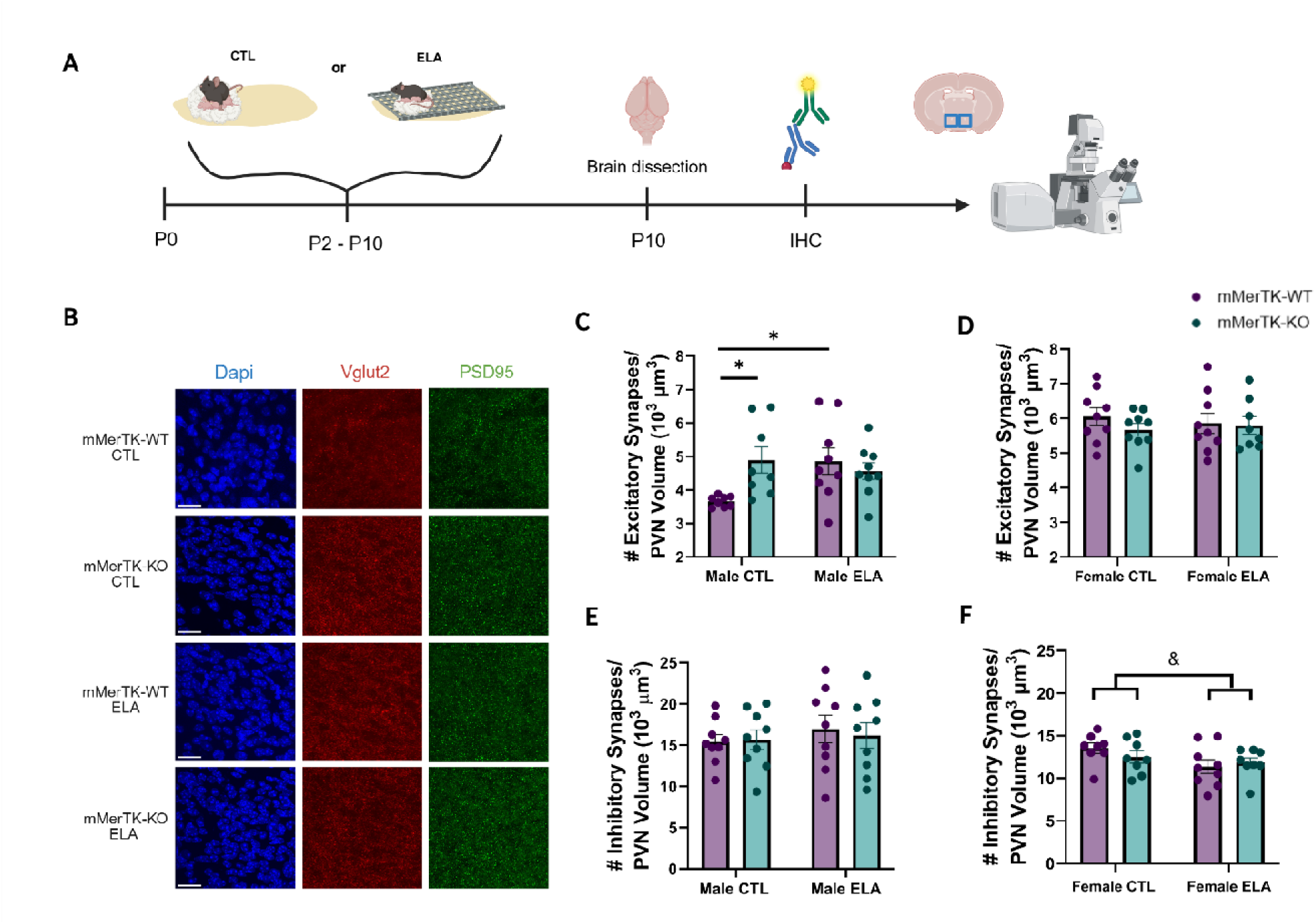
Microglial MerTK-KO in control males mimics the ELA-induced increase in excitatory synapses in the PVN of males. (A) Timeline for IHC experiments: Male and female mice, randomly assigned to CTL or ELA conditions, were perfused on P10. IHC was then performed and excitatory and inhibitory synapses onto mpd PVN neurons were quantified. The graphic was created with https://BioRender.com (B) Representative images showing DAPI, vGlut2 and PSD95 staining in the mpd PVN of all four experimental groups in males (mMerTK-WT CTL, mMerTK-KO CTL, mMerTK-WT ELA, mMerTK-KO ELA), Scale bar=20 μm. (C) Deletion of microglial MerTK *in vivo* increases the number of excitatory synapses onto mpd PVN neurons in control males to the level of ELA males, but causes no further increase in ELA males. (D) There is no effect of mMerTK-KO nor ELA on excitatory synapses in the mpd PVN of females, although they are higher at baseline than males. (E) Neither ELA nor mMerTK-KO alters inhibitory synapse number in the mpd PVN of males. (F) ELA tends to decrease the number of inhibitory synapses in females. Data are means ± SEMs; \**p*<0.05, post hoc Šídák’s test, following up ELA x genotype interaction; ^&^*p =* 0.05, trend for main effect of ELA.

### 3.2 ELA increases the neuroendocrine response to stress in males

The mpd PVN is densely populated with CRH-expressing neurons, which are essential for the neuroendocrine response to stress by the HPA axis. ELA has previously been shown to alter the hormonal response to stress in adulthood in males (Bolton et al., 2022), but the stress response in ELA females is less well-characterized. Therefore, restraint-stress testing was performed on adult male and female mice of each genotype, reared in either CTL or ELA conditions. Acute stress response was assessed by measuring corticosterone (CORT) levels across several timepoints, and a surrogate measure of the chronic stress response was assessed by measuring the weights of adrenal glands (Ulrich-Lai et al., 2006, Harper & Austad, 2000; Figure 4A). At baseline, the release of CORT was found to be increased by ELA in mMerTK-WT mice of both sexes, as previously (Rice et al., 2008), although mMerTK-KO mice did not differ (trend for Genotype x ELA interaction, *F*_1, 52_= 2.396, *p*=0.1; 3-way ANOVA; *p*<0.05; post hoc Šídák’s test; Figure 4F-G). ELA increased the adrenal weights of mMerTK-WT males (significant Genotype x ELA interaction, *F*_1, 34_= 4.797, *p*=0.04; 2-way ANOVA, *p*=0.08; post hoc Šídák’s test; Figure 4H) and mMerTK-KO males and females (*p*<0.05; post hoc Šídák’s test), but not mMerTK-WT females, whose adrenal weights were already higher at baseline (significant main effect of Sex, *F*_1, 69_= 32.64, *p*<0.0001; trend for LBN x Sex interaction, *F*_1, 69_= 2.265, *p*=0.1; 3-way ANOVA). Additionally, mMerTK-KO increased adrenal weights in both ELA males (p=0.08; post hoc Šídák’s test; Figure 4H) and females (trend for Genotype x ELA interaction, *F*_1, 33_= 2.650, *p*=0.1; 2-way ANOVA,\**p*=0.08; post hoc Šídák’s test; Figure 4I). However, ELA elevated the CORT response to acute restraint stress in males (significant main effect of ELA, *F*_1, 30_= 11.15, *p*=0.002; 2-way ANOVA; Figure 4B), but had no effect in females, which were already elevated at baseline (significant main effect of Sex, *F*_1, 58_= 74.78, *p*<0.0001; trend for ELA x sex interaction, *F*_1, 58_= 2.410, *p*=0.1; 3-way ANOVA; Figure 4E). Surprisingly, there was also no effect of the mMerTK-KO on CORT levels or the weight of adrenal glands in either sex (trend for ELA x Sex interaction, *F*_1, 69_= 2.265, *p*=0.1; 3-way ANOVA; Figure 4H-I).

**Figure 4.**
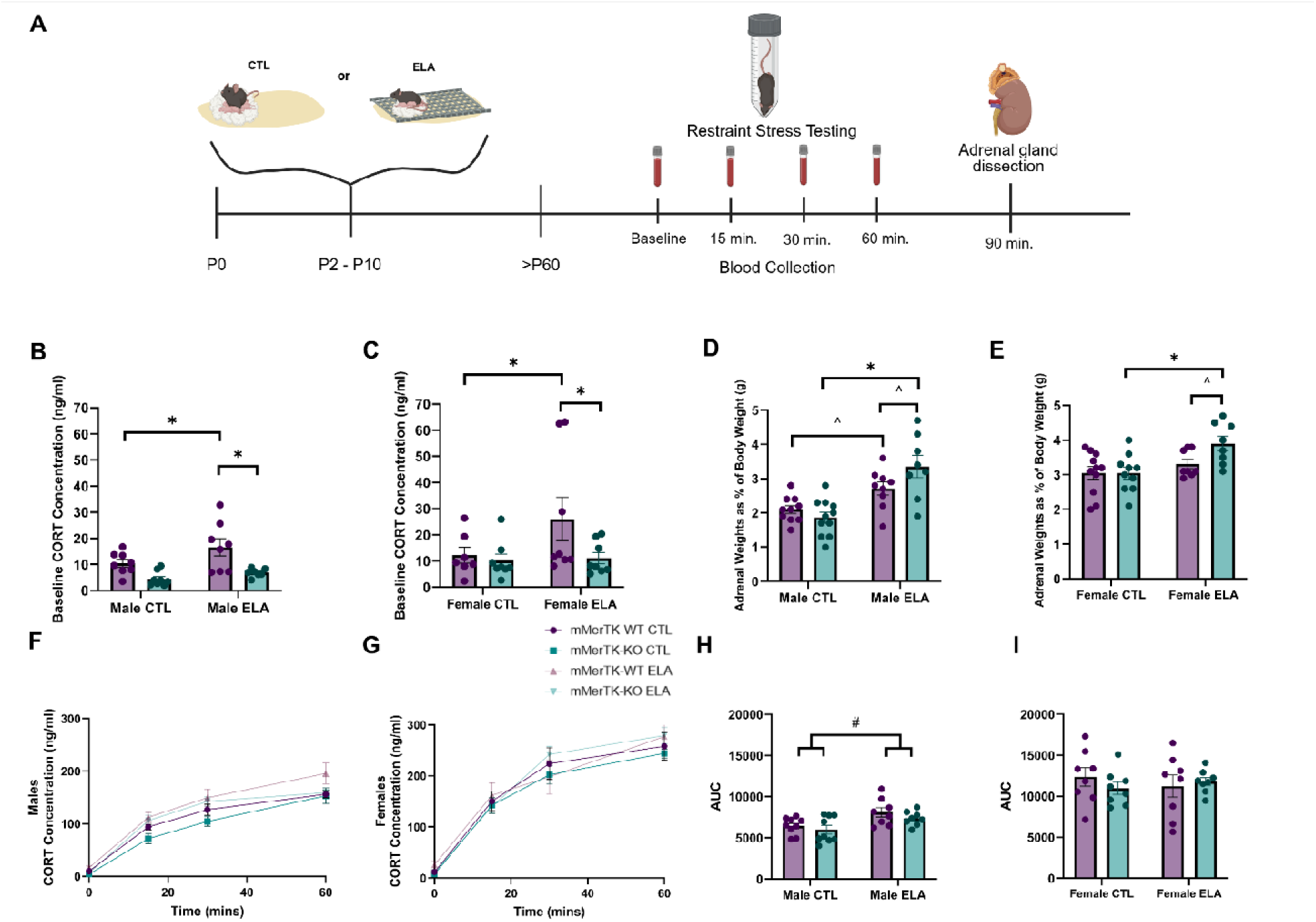
The neuroendocrine response to stress is altered primarily by ELA in males. (A) Schematic of the paradigm to assess the acute and chronic neuroendocrine response to stress: Male and female mice of both genotypes, reared in CTL or ELA conditions, were subjected to acute restraint stress, in which they were placed in a 50-mL conical tube for 90 min. Blood was collected for stress hormone analysis at baseline, 15 min., 30 min., and 60 min. Adrenal glands were collected and weighed at 90 min. The diagram was created with https://BioRender.com (B-C) ELA increases baseline CORT level in mMerTK-WT, but not mMerTK-KO, males (B) and females (C). (D) Adrenal weights of adult ELA males are higher in both mMertK-WT and mMerTK-KO mice, and mMerTK-KO males are higher than mMerTK-WT males within the ELA group. (E) Adrenal weights of ELA females are higher in mMerTK-KO, but not mMerTK-WT mice, and mMerTK-KO females are higher than mMerTK-WT females within the ELA group. (F-G) Serum CORT concentrations of all four experimental groups (mMerTK-WT CTL, mMerTK-KO CTL, mMerTK-WT ELA, mMerTK-KO ELA) over 60 min. of restraint stress in males (F) and (G) females. (H-I) The CORT response over 60 min., as measured by area under the curve (AUC) from F-G, is significantly elevated by ELA in males (H) but not females (I). Data are means ± SEMs. ^^^*p*=0.08, \**p*<0.05, post hoc Sidak’s test, following up ELA x Genotype interaction. ^#^*p*<0.05, main effect of ELA.

### 3.3 Aberrant threat-response behavior provoked by ELA is recapitulated by microglial MerTK-KO in control males

The long-lasting effects of ELA on the stress response have also been found to reach beyond hormonal changes, leading to aberrant behavioral responses to threat (Bolton et al., 2022). Therefore, in order to further probe the functional significance of MerTK in ELA-induced alterations to threat-response behavior, we implemented the looming-shadow threat task (Figure 5A-B). Consistent with previous findings (Short et al., 2021), ELA augmented threat-response behavior in male mice, as measured by increased escape behavior (significant Genotype x ELA interaction, *F*_1, 38_ = 4.475, *p*=0.04; 2-way ANOVA; *p*<0.05; post hoc Šídák’s test; Figure 5C) and a decreased latency to escape in mMerTK-WT males (trend for Genotype x ELA interaction, *F*_1, 31_ = 3.194, *p*=0.08; 2-way ANOVA; *p* = 0.06; post hoc Šídák’s test; Figure 5E). This was mimicked by mMerTK-KO in control males (*p*<0.05; post hoc Šídák’s test; Figure 5C). Notably, there was no further increase in escape behavior due to mMerTK-KO in the ELA group, again indicating that this pathway is already occluded by ELA (Figure 5C). There was no significant effect of ELA or mMerTK-KO on freezing behavior in males or females *(p*>0.1; 3-way ANOVA; Supplemental Figure 2A). There was also no effect of ELA or the mMerTK-KO on escape behavior (Genotype x ELA x Sex, *F*_1, 67_= 5.729, *p*=0.02; 3-way ANOVA; Figure 5D) or the latency to escape for females (significant interaction of Genotype x ELA x Sex, *F*_1, 63_= 4.371, *p*=0.04; 3-way ANOVA; Figure 5E-F).

**Figure 5.**
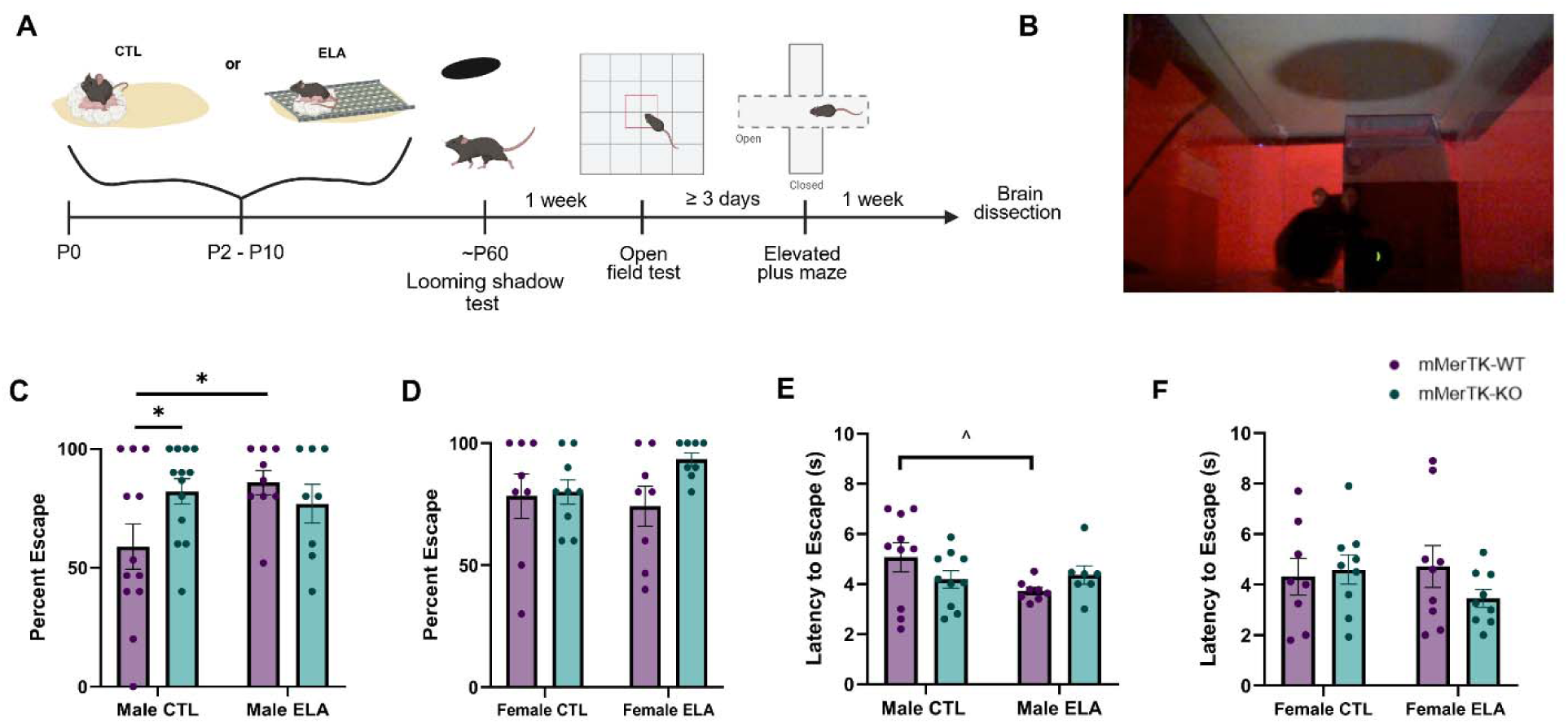
Deleting microglial MerTK recapitulates the effect of ELA on threat-response behavior in males. (A) Diagram of timeline for behavioral experiments: Adult (∼P60) male and female mice of both genotypes, reared in CTL or ELA conditions, were run through a behavioral battery consisting of the looming-shadow threat task, open-field test and elevated-plus maze. The graphic was created with https://BioRender.com. (B) Representative image of the LST: subjects were presented with the looming-shadow stimulus 5 times and responses were recorded as escape, freeze or none. (C) mMertk-KO increases escape behavior in the LST in control males to the level of ELA but does not further increase escape in ELA males. (D) Escape behavior is not significantly altered by the mMerTK-KO or ELA in females. (E-F) There was a trend for ELA to decrease the latency to escape in mMerTK-WT males (E), but not females (F). Data are means ± SEMs. *p<0.05; ^^^p = 0.06; post hoc Sidak’s test, following up ELA x genotype interaction.

### 3.4 ELA, but not mMerTK-KO, alters anxiety-like behavior in females more than males

Lastly, to ascertain if the MerTK pathway underlies any enduring effects of ELA on anxiety-like behavior, we performed the OFT and EPM in adulthood (Figure 6A). As expected, there was no difference observed in the time spent in the center of the open field arena in male mice (significant ELA x Sex interaction, *F*_1, 69_= 6.988, *p*=0.001; 3-way ANOVA; Figure 6A, E). However, ELA decreased the time spent in the center of the open field in female mice, suggesting increased anxiety-like behavior (significant main effect of ELA, *F*_1, 34_ = 5.945 *p*=0.02; 2-way ANOVA; Figure 6B). Surprisingly, ELA increased the time spent in the open arms of the EPM in mMerTK-WT mice of both sexes (trend for Genotype x ELA interaction, *F*_1, 68_= 3.457, *p*=0.07; 3-way ANOVA,*p<0.05; post hoc Šídák’s test; Figure 6F) and decreased the time spent in the closed arms of the EPM in both sexes (trend for Genotype x ELA interaction, *F*_1, 70_= 3.708, *p*=0.06; 3-way ANOVA,*p<0.05; post hoc Šídák’s test; Supplemental Figure 3B), suggesting a decrease in anxiety-like behavior, or increase in risk-taking behavior, in that task in both sexes. The mMerTK-KO did not alter anxiety-like behavior in either task in males or females (Figure 6A, B, E, F). Importantly, there was no effect of ELA nor mMerTK-KO on total distance travelled in the OFT (*p*>0.5; 3-way ANOVA; Figure 6C-D) or EPM (*p*>0.4; 3-way ANOVA; Figure 6G-H) in either sex, suggesting that ELA did not induce general changes in locomotion or hyperactivity

**Figure 6:**
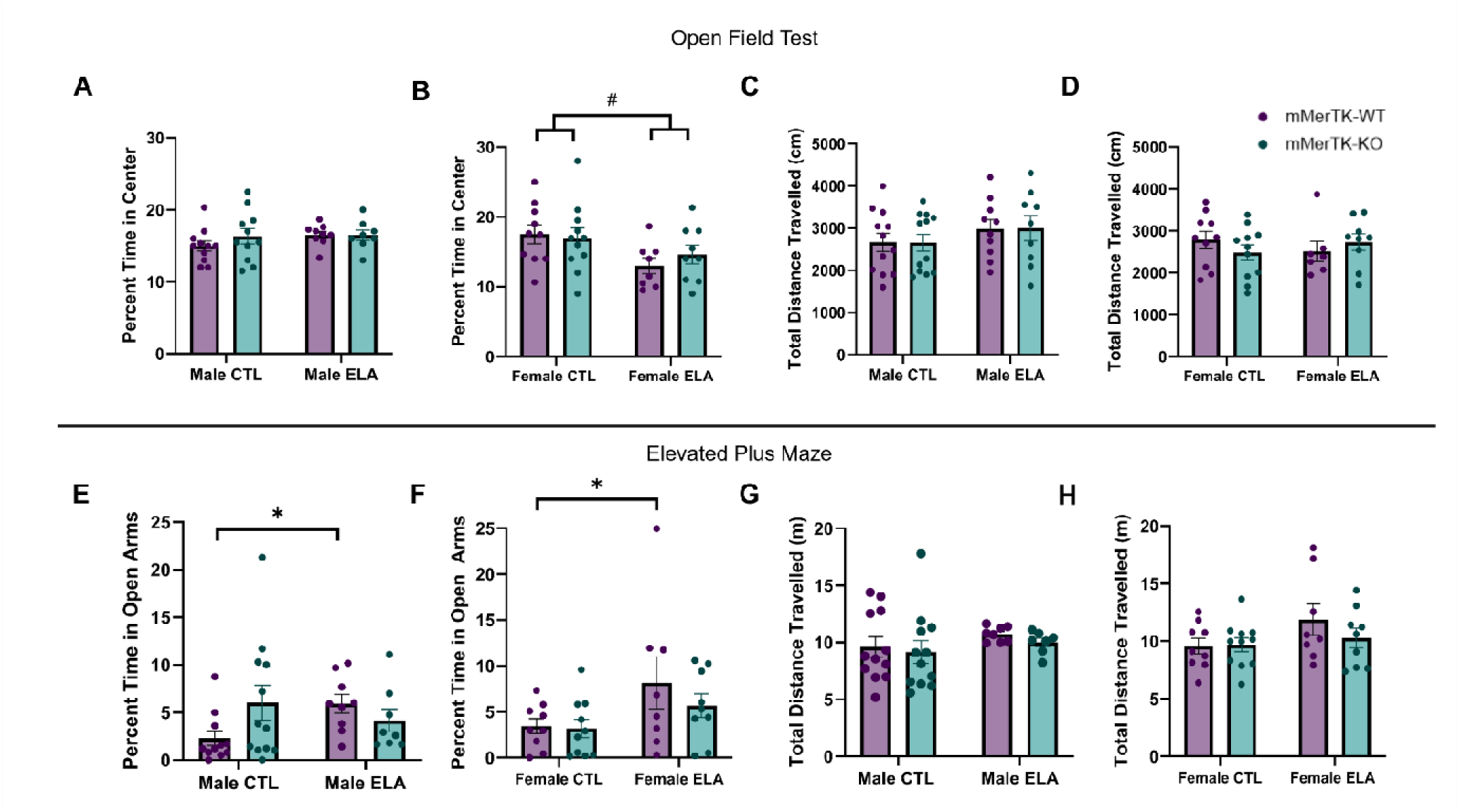
ELA, but not microglial MerTK-KO, alters anxiety-like behavior more in females. (A) Percent time in the center of the open-field test does not differ by ELA or genotype in males. (B) ELA decreases the percent time spent in the center of the OFT in female mice. (C-D) ELA does not alter total distance travelled in the OFT in males (C) or females (D). (E-F) ELA increases the percent time spent in the open arms in mMerTK-WT males (E) and females (F). (G-H) Total distance travelled in the EPM does not differ between control and ELA conditions in males (G) or females (H). Data are means ± SEMs. \**p*<0.05, post hoc Sidak’s test, following up ELA x Genotype interaction. ^#^*p*<0.05, main effect of ELA.

## 4. Discussion

Here, we demonstrate a critical role for the microglial phagocytic receptor, MerTK, in the development of excitatory synapses onto stress-sensitive neurons in the hypothalamus and threat-response behavior in adulthood. We have previously characterized the effects of ELA on microglial function and stress-circuit maturation in the developing PVN and identified MerTK as a potential underlying factor (Bolton et al., 2022). However, the evidence supporting this mechanism was primarily *in vitro* and lacking cell-type specificity. Therefore, here we employed a microglia-specific MerTK-KO mouse line in order to further elucidate the extent to which this molecular pathway underlies the effects of ELA at both the cellular and behavioral level.

Deleting microglial MerTK had a significant impact on synapse development in the early postnatal period of males. The mMerTK-KO in control male mice resulted in an increase in the number of excitatory synapses in the mpd PVN, to the level of ELA males. Importantly, the mMerTK-KO did not provoke the same effects in the ELA group, demonstrating that this pathway is already inhibited by ELA (i.e., an occlusion effect). This is congruent with our previous findings showing that MerTK expression is decreased by ELA in the immature male PVN (Bolton et al., 2022). However, ELA did not provoke a significant increase in excitatory synapses in females, although their numbers were already increased in controls. Perhaps because of this baseline sex difference and the decreased level of MerTK expressed in control females, mMerTK-KO did not cause any significant change in synapses. Thus, these results indicate that microglial MerTK is critical for the normal maturation of stress-related circuitry in the PVN of males, but not females, and likely underlies the sex-specific aberrant synapse development induced by ELA.

Escape behavior in the looming-shadow threat test has been shown to be dependent on increased CRH+ neuron activity in the PVN (Daviu, Füzesi, Rosenegger, Rasiah, et al., 2020). In line with previous work (Short et al., 2021), we found that ELA significantly increased escape behavior in males. Importantly, we also observed an augmented escape response due to the mMerTK-KO in male control mice, but not ELA mice. These results further support the importance of microglial MerTK in not only the development of synapses onto stress-sensitive neurons but also their functional output, such as the behavioral response to a threat in adulthood in males. Similar to the analysis of excitatory synapses in the PVN, we did not find an effect of ELA or mMerTK-KO on escape behavior in females, indicating this pathway may play less of a role in females than in males.

Due to the role of the PVN in regulating the neuroendocrine response to stress, we also analyzed hormonal responses to acute stress testing in adulthood. We found that ELA amplified CORT responses to stress in males and increased adrenal weights in both sexes, indicating it alters both the acute and chronic stress response in mice, as previously (Bolton et al., 2022). Surprisingly, we observed no effect of the mMerTK-KO in males or females. However, neuroendocrine changes may be affected by peripheral factors in addition to brain-derived factors. It is important to note that the transgenic approach we employed here also deletes MerTK in CX3CR1+ peripheral macrophages/monocytes in the rest of the body. This could have resulted in some of the unexpected changes in adrenal weights and the peripheral CORT response to acute stress. However, there is currently no evidence for effects of these CX3CR1+ peripheral cells on localized synapse numbers in the PVN during development. Using tamoxifen-inducible Cre mice (e.g., P2RY12-CreER) is an alternative strategy that would result in more specificity to microglia vs. peripheral macrophages, but our interest in sex differences makes the neonatal administration of tamoxifen, an estrogen receptor modulator, problematic. Thus, we propose that the approach we have taken here to conditionally knockout MerTK in microglia in transgenic mouse pups (CX3CR1-Cre^+^::Mertk^fl/fl^) is the best option currently available.

Despite their apparent resilience to the effects of ELA in regard to threat-response behavior, which may be due to their increased stress-responsivity at baseline, we did observe an effect of ELA on anxiety-like behavior in females. In the open-field test, ELA decreased the time spent in the center of the OFT for female mice, which may indicate increased anxiety-like behavior. Perhaps more intriguingly, ELA increased the time spent in the open arms of the EPM in both male and female mMertk-WT mice, perhaps indicating decreased anxiety-like behavior or increased risk-taking behavior. It is unclear why the two tests of anxiety-like behavior produced contradictory results in females, although this has type of disagreement been reported before in the literature, and may be due to the different contexts of each test (i.e., the EPM is more of a novel, and potentially more threatening, environment).

There is widespread support in the literature for the greater vulnerability of males across species to the effects of ELA and other perinatal challenges, perhaps due to their brains undergoing the complex process of masculinization by gonadal hormones during this same sensitive period. While the focus has been on male-specific vulnerability, and a majority of studies using this ELA model have examined only male animals, it is noteworthy that in most of our data there are robust baseline sex differences, with females being more stress-reactive overall, and that ELA actually appear to make males more “female-like”. Evolutionarily speaking, it may be important for masculinization to render males less stress-responsive than females in order to foster mate-seeking and territory-guarding behavior. Thus, it is possible that ELA may interfere with masculinization of the brain, making males more stress-responsive, whereas females appear resilient to ELA because they are already functionally “maxed-out” for stress responsivity, which is potentially more adaptive for their maternal role. The ultimate basis of the baseline sex difference and the response to ELA is the focus of ongoing work in our lab.

This research reveals, for the first time, sex- and cell type-specific mechanisms regulating the synaptic pruning of developing stress circuits, and the normal expression of stress-/threat-response behavior in adulthood. Importantly, these findings provide the foundation for developing more targeted and personalized preventative and therapeutic strategies for neuropsychiatric disorders like depression in at-risk children.

## Acknowledgments

The authors would like to thank the Division of Animal Resources at Georgia State University for providing exceptional care to our animals.

## Appendix A. Supplementary data

**Supplemental Figure 1.**
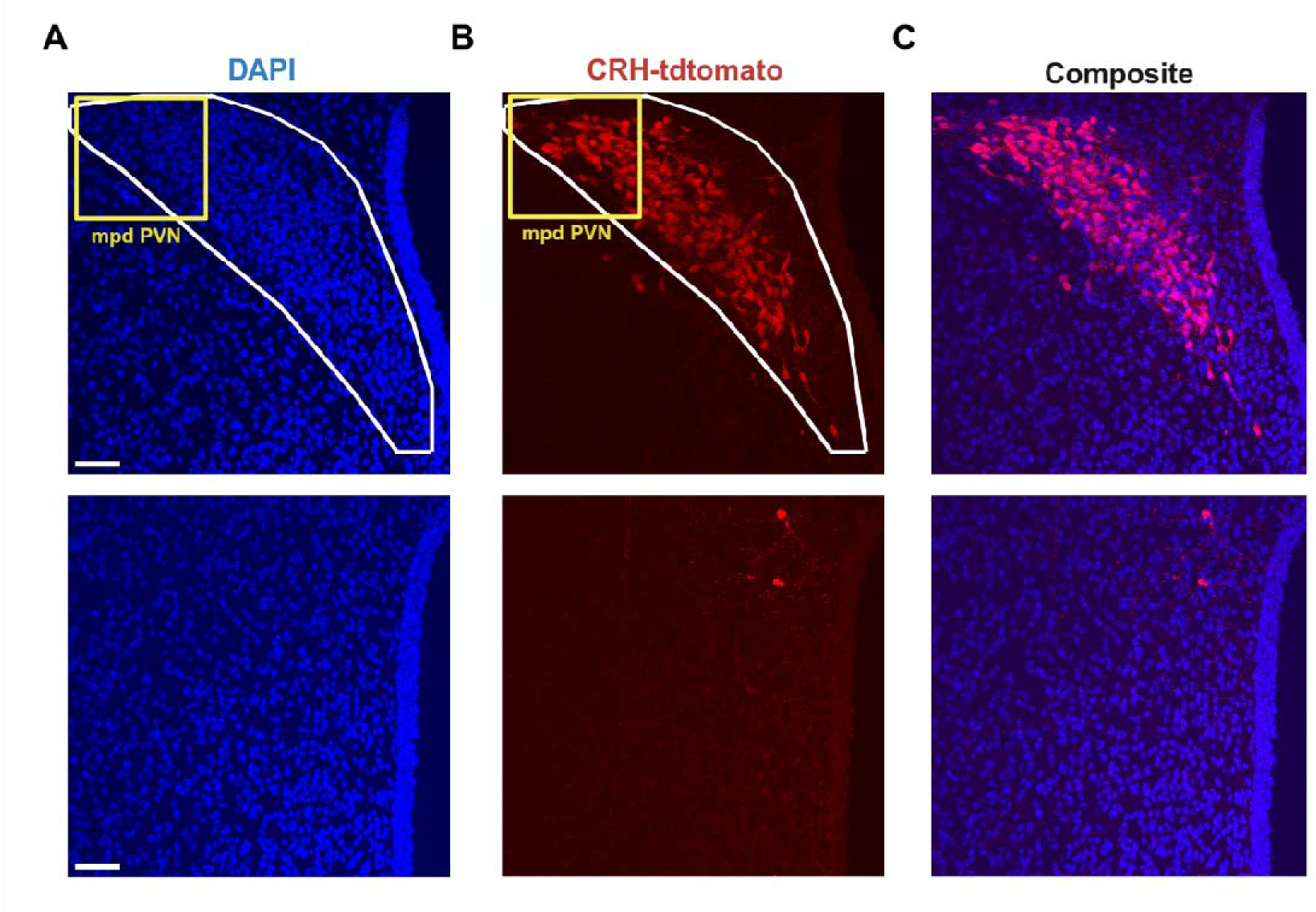
Representative images of overlap between (A) DAPI and (B) CRH-tdTomato+ neurons in the PVN (top) and posterior to the PVN (bottom) at P8.

**Supplemental Figure 2.**
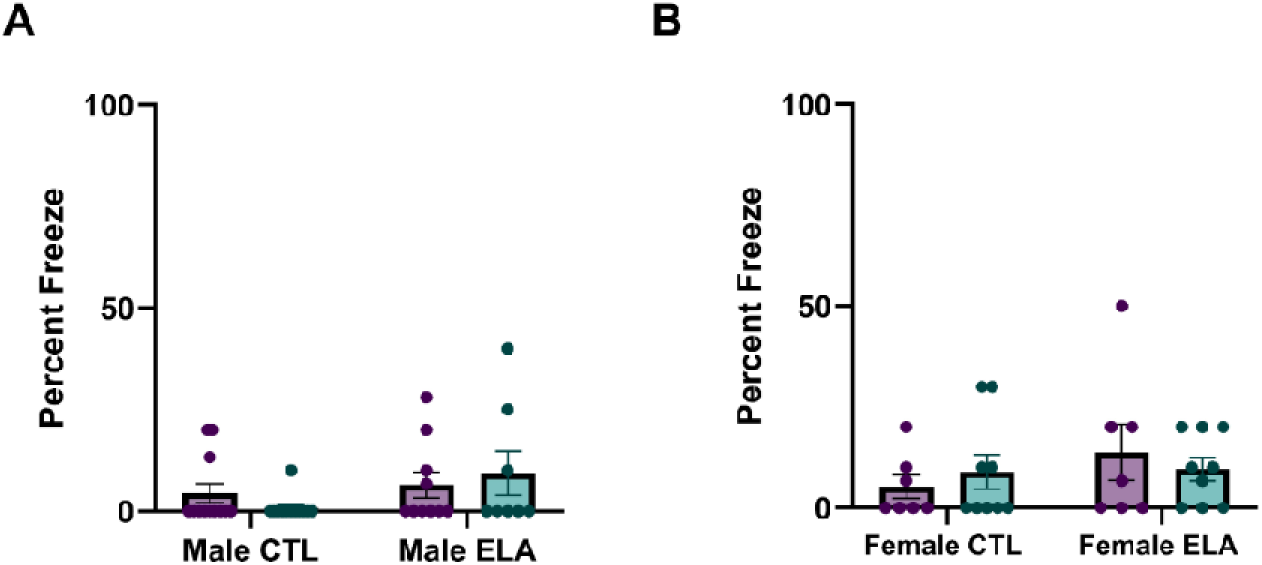
A-B) There was no significant difference in freezing behavior in the looming-shadow threat task in males (A) or females (B).

**Supplemental Figure 3.**
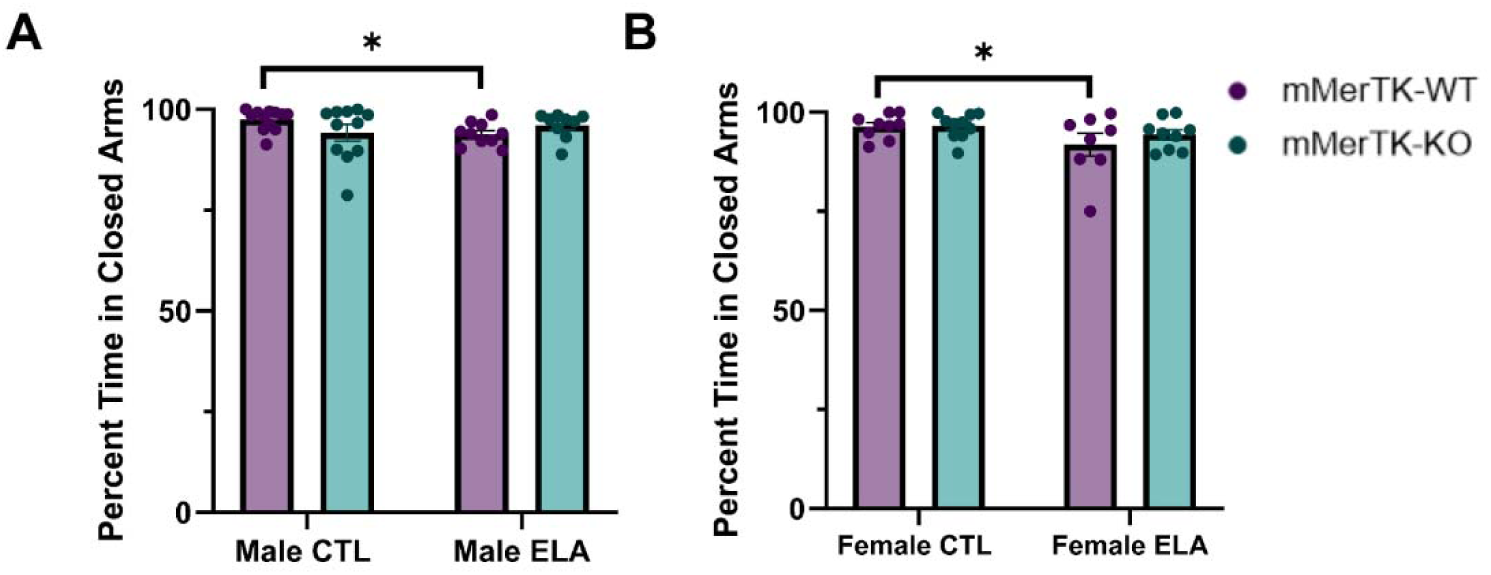
(A-B) ELA decreased the time spent in the closed arms of the EPM in mMerTK-WT mice of both sexes.

## Data availability

Data will be made available on request.

## References

1. Abiega, O., Beccari, S., Diaz-Aparicio, I., Nadjar, A., Layé, S., Leyrolle, Q., Gómez-Nicola, D., Domercq, M., Pérez-Samartín, A., Sánchez-Zafra, V., Paris, I., Valero, J., Savage, J. C., Hui, C. W., Tremblay, M. È., Deudero, J. J. P., Brewster, A. L., Anderson, A. E., Zaldumbide, L., … Sierra, A. (2016). Neuronal Hyperactivity Disturbs ATP Microgradients, Impairs Microglial Motility, and Reduces Phagocytic Receptor Expression Triggering Apoptosis/Microglial Phagocytosis Uncoupling. PLoS Biology, 14(5), 1–48. 10.1371/journal.pbio.1002466

2. Ahmed, S., Polis, B., Jamwal, S., Sanganahalli, B. G., Kaswan, Z. M., Islam, R., Kim, D., Bowers, C., Giuliano, L., Biederer, T., Hyder, F., & Kaffman, A. (2024). Transient Impairment in Microglial Function Causes Sex-Specific Deficits in Synaptic and Hippocampal Function in Mice Exposed to Early Adversity. BioRxiv, 2, 2024.02.14.580284. 10.1101/2024.02.14.580284

3. Amateau, S. K., & McCarthy, M. M. (2002). A Novel Mechanism of Dendritic Spine Plasticity Involving Estradiol Induction of Prostaglandin-E2. Journal of Neuroscience, 22(19), 8586– 8596. 10.1523/JNEUROSCI.22-19-08586.2002

4. Amateau, S. K., & McCarthy, M. M. (2004). Induction of PGE2 by estradiol mediates developmental masculinization of sex behavior. Nature Neuroscience, 7(6), 643–650. 10.1038/NN1254

5. Bick, J., & Nelson, C. A. (2016). Early Adverse Experiences and the Developing Brain. Neuropsychopharmacology, 41(1), 177–196. 10.1038/npp.2015.252

6. Bilbo, S. D., & Schwarz, J. M. (2012). The immune system and developmental programming of brain and behavior. 33(3). 10.1016/j.yfrne.2012.08.006

7. Bolton, J. L., Molet, J., Regev, L., Chen, Y., Rismanchi, N., Haddad, E., Yang, D. Z., Obenaus, A., & Baram, T. Z. (2018). Anhedonia Following Early-Life Adversity Involves Aberrant Interaction of Reward and Anxiety Circuits and Is Reversed by Partial Silencing of Amygdala Corticotropin-Releasing Hormone Gene. Biological Psychiatry, 83(2), 137–147. 10.1016/j.biopsych.2017.08.023

8. Bolton, J. L., Short, A. K., Othy, S., Kooiker, C. L., Shao, M., Gunn, B. G., Beck, J., Bai, X., Law, S. M., Savage, J. C., Lambert, J. J., Belelli, D., Tremblay, M. È., Cahalan, M. D., & Baram, T. Z. (2022a). Early stress-induced impaired microglial pruning of excitatory synapses on immature CRH-expressing neurons provokes aberrant adult stress responses. Cell Reports, 38(13). 10.1016/j.celrep.2022.110600

9. Chung, W. S., Clarke, L. E., Wang, G. X., Stafford, B. K., Sher, A., Chakraborty, C., Joung, J., Foo, L. C., Thompson, A., Chen, C., Smith, S. J., & Barres, B. A. (2013). Astrocytes mediate synapse elimination through MEGF10 and MERTK pathways. Nature, 504(7480), 394–400. 10.1038/nature12776

10. Chung, W.-S., Clarke, L. E., Wang, G. X., Stafford, B. K., Sher, A., Chakraborty, C., Joung, J., Foo, L. C., Thompson, A., Chen, C., Smith, S. J., & Barres, B. A. (2013). Astrocytes mediate synapse elimination through {MEGF10} and {MERTK} pathways. 504(7480), 394–400. 10.1038/nature12776

11. Colle, R., Segawa, T., Chupin, M., Tran Dong, M. N. T. K., Hardy, P., Falissard, B., Colliot, O., Ducreux, D., & Corruble, E. (2017). Early life adversity is associated with a smaller hippocampus in male but not female depressed in-patients: A case-control study. BMC Psychiatry, 17(1), 1–8. 10.1186/S12888-017-1233-2/FIGURES/2

12. Danese, A., & McEwen, B. S. (2012). Adverse childhood experiences, allostasis, allostatic load, and age-related disease. Physiology & Behavior, 106(1), 29–39. 10.1016/j.physbeh.2011.08.019

13. Davis, E. C., Popper, P., & Gorski, R. A. (1996). The role of apoptosis in sexual differentiation of the rat sexually dimorphic nucleus of the preoptic area. Brain Research, 734(1–2), 10–18. 10.1016/0006-8993(96)00298-3

14. Daviu, N., Füzesi, T., Rosenegger, D. G., Peringod, G., Simone, K., & Bains, J. S. (2020). Visual-looming Shadow Task with in-vivo Calcium Activity Monitoring to Assess Defensive Behaviors in Mice. Bio-Protocol, 10(22). 10.21769/BioProtoc.3826

15. Daviu, N., Füzesi, T., Rosenegger, D. G., Rasiah, N. P., Sterley, T. L., Peringod, G., & Bains, J. S. (2020a). Paraventricular nucleus CRH neurons encode stress controllability and regulate defensive behavior selection. Nature Neuroscience, 23(3), 398–410. 10.1038/s41593-020-0591-0

16. Diaz-Aparicio, I., Paris, I., Sierra-Torre, V., Plaza-Zabala, A., Rodríguez-Iglesias, N., Márquez-Ropero, M., Beccari, S., Huguet, P., Abiega, O., Alberdi, E., Matute, C., Bernales, I., Schulz, A., Otrokocsi, L., Sperlagh, B., Happonen, K. E., Lemke, G., Maletic-Savatic, M., Valero, J., & Sierra, A. (2020). Microglia Actively Remodel Adult Hippocampal Neurogenesis through the Phagocytosis Secretome. The Journal of Neurosciencel: The Official Journal of the Society for Neuroscience, 40(7), 1453–1482. 10.1523/JNEUROSCI.0993-19.2019

17. Fabricius, K., Wörtwein, G., & Pakkenberg, B. (2008). The impact of maternal separation on adult mouse behaviour and on the total neuron number in the mouse hippocampus. Brain Structure and Function, 212(5), 403–416. 10.1007/s00429-007-0169-6

18. Faust, T. E., Gunner, G., & Schafer, D. P. (2021). Mechanisms governing activity-dependent synaptic pruning in the developing mammalian CNS. Nature Reviews Neuroscience 2021 22:11, 22(11), 657–673. 10.1038/s41583-021-00507-y

19. Fourgeaud, L., Través, P. G., Tufail, Y., Leal-bailey, H., Lew, E. D., Burrola, P. G., Nimmerjahn, A., Lemke, G., Callaway, P., Zagórska, A., & Carla, V. (2016). TAM receptors regulate multiple features of microglial physiology. Nature, 532(7598), 240–244. 10.1038/nature17630

20. Garvin, M. M., & Bolton, J. L. (2022). Sex-specific behavioral outcomes of early-life adversity and emerging microglia-dependent mechanisms. Frontiers in Behavioral Neuroscience, 16, 1013865. 10.3389/fnbeh.2022.1013865

21. Gosselin, D., Skola, D., Coufal, N. G., Holtman, I. R., Schlachetzki, J. C. M., Sajti, E., Jaeger, B. N., O’Connor, C., Fitzpatrick, C., Pasillas, M. P., Pena, M., Adair, A., Gonda, D. G., Levy, M. L., Ransohoff, R. M., Gage, F. H., & Glass, C. K. (2017). An environment-dependent transcriptional network specifies human microglia identity. Science. 10.1126/science.aal3222

22. Harper, J. M., & Austad, S. N. (2000). Fecal glucocorticoids: A noninvasive method of measuring adrenal activity in wild and captive rodents. Physiological and Biochemical Zoology, 73(1), 12–22. 10.1086/316721

23. Heim, C., & Binder, E. B. (2012). Current research trends in early life stress and depression: Review of human studies on sensitive periods, gene–environment interactions, and epigenetics. Experimental Neurology, 233(1), 102–111. 10.1016/j.expneurol.2011.10.032

24. Hoogland, M., & Ploeger, A. (2022). Two Different Mismatches: Integrating the Developmental and the Evolutionary-Mismatch Hypothesis. Perspectives on Psychological Science, 17(6), 1737–1745. 10.1177/17456916221078318/ASSET/IMAGES/LARGE/10.1177_17456916 221078318-FIG1.JPEG

25. Lenz, K. M., Nugent, B. M., Haliyur, R., & M, M. M. (2013). Microglia are essential to masculinization of brain and behavior. 33(7), 2761–2772. 10.1523/JNEUROSCI.1268-12.2013

26. Malcon, L. M. C., Wearick-Silva, L. E., Zaparte, A., Orso, R., Luft, C., Tractenberg, S. G., Donadio, M. V. F., de Oliveira, J. R., & Grassi-Oliveira, R. (2020). Maternal separation induces long-term oxidative stress alterations and increases anxiety-like behavior of male Balb/cJ mice. Experimental Brain Research, 238(9), 2097–2107. 10.1007/s00221-020-05859-y

27. McCarthy, M. M. (2016). Sex differences in the developing brain as a source of inherent risk. Dialogues in Clinical Neuroscience, 18(4), 361–372. 10.31887/DCNS.2016.18.4/MMCCARTHY

28. McCarthy, M. M., Amateau, S. K., & Mong, J. A. (2002). Steroid Modulation of Astrocytes in the Neonatal Brain: Implications for Adult Reproductive Function. Biology of Reproduction, 67(3), 691–698. 10.1095/BIOLREPROD.102.003251

29. McCarthy, M. M., Todd, B. J., & Amatea, S. K. (2003). Estradiol Modulation of Astrocytes and the Establishment of Sex Differences in the Brain. Annals of the New York Academy of Sciences, 1007(1), 283–297. 10.1196/ANNALS.1286.027

30. McCarthy, M. M., & Wright, C. L. (2017). Convergence of Sex Differences and the Neuroimmune System in Autism Spectrum Disorder. Biological Psychiatry, 81(5), 402–410. 10.1016/J.BIOPSYCH.2016.10.004

31. McLaughlin, K. A., Weissman, D., & Bitrán, D. (2019). Childhood Adversity and Neural Development: A Systematic Review. Annual Review of Developmental Psychology, 1(1), 277–312. 10.1146/annurev-devpsych-121318-084950

32. Mong, J. A., & McCarthy, M. M. (2002). Ontogeny of sexually dimorphic astrocytes in the neonatal rat arcuate. Developmental Brain Research, 139(2), 151–158. 10.1016/S0165-3806(02)00541-2

33. Mroue-Ruiz, F. H., Garvin, M., Ouellette, L., Sequeira, M. K., Lichtenstein, H., Kar, U., & Bolton, J. L. (2024). Limited Bedding and Nesting as a Model for Early-Life Adversity in Mice. Journal of Visualized Experiments, 2024(209). 10.3791/66879

34. Paolicelli, R. C., Bolasco, G., Pagani, F., Maggi, L., Scianni, M., Panzanelli, P., Giustetto, M., Ferreira, T. A., Guiducci, E., Dumas, L., Ragozzino, D., & Gross, C. T. (2011). Synaptic Pruning by Microglia Is Necessary for Normal Brain Development. Science, 333(6048), 1456–1458.

35. Pickett, L. A., VanRyzin, J. W., Marquardt, A. E., & McCarthy, M. M. (2023). Microglia phagocytosis mediates the volume and function of the rat sexually dimorphic nucleus of the preoptic area. Proceedings of the National Academy of Sciences of the United States of America, 120(10). 10.1073/PNAS.2212646120

36. Rice, C. J., Sandman, C. A., Lenjavi, M. R., & Baram, T. Z. (2008). A novel mouse model for acute and long-lasting consequences of early life stress. Endocrinology, 149(10), 4892– 4900. 10.1210/en.2008-0633

37. Schafer, D., Lehrman, E., Kautzman, A., Koyama, R., Mardinly, A., Yamasaki, R., Ransohoff, R., Greenberg, M., Barres, B., & Stevens, B. (2012). Microglia sculpt postnatal neuronal circuits in an activivty and complement-dependent manner. Neuron, 74(4), 691–705. 10.1016/j.neuron.2012.03.026.Microglia

38. Schwarz, J. M., & Bilbo, S. D. (2012). Sex, glia, and development: Interactions in health and disease. Hormones and Behavior.

39. Schwarz, J. M., Sholar, P. W., & Bilbo, S. D. (2012). Sex differences in microglial colonization of the developing rat brain. Journal of Neurochemistry, 948–963.

40. Scott-Hewitt, N., Perrucci, F., Morini, R., Erreni, M., Mahoney, M., Witkowska, A., Carey, A., Faggiani, E., Schuetz, L. T., Mason, S., Tamborini, M., Bizzotto, M., Passoni, L., Filipello, F., Jahn, R., Stevens, B., & Matteoli, M. (2020). Local externalization of phosphatidylserine mediates developmental synaptic pruning by microglia. The EMBO Journal, 39(16), e105380. 10.15252/embj.2020105380

41. Seillier, A., & Giuffrida, A. (2017). Anxiety does not contribute to social withdrawal in the subchronic phencyclidine rat model of schizophrenia. Behavioural Pharmacology, 28(7), 512–520. 10.1097/FBP.0000000000000325

42. Short, A. K., Wilcox, C., Chen, Y., Pham, A. L., Birnie, M. T., Bolton, J. L., Mortazavi, A., & Baram, T. Z. (2021). Single-Cell transcriptional changes in hypothalamic CRH-expressing neurons after early-life adversity inform enduring alterations in responses to stress. BioRxiv, 2021.08.31.458231. 10.1101/2021.08.31.458231

43. Simon, P., Dupuis, R., & Costentin, J. (1994). Thigmotaxis as an index of anxiety in mice. Influence of dopaminergic transmissions. Behavioural Brain Research, 61(1), 59–64. 10.1016/0166-4328(94)90008-6

44. Sorge, R. E., Mapplebeck, J. C. S., Rosen, S., Beggs, S., Taves, S., Alexander, J. K., Martin, L. J., Austin, J. S., Sotocinal, S. G., Chen, D., Yang, M., Shi, X. Q., Huang, H., Pillon, N. J., Bilan, P. J., Tu, Y., Klip, A., Ji, R. R., Zhang, J., … Mogil, J. S. (2015). Different immune cells mediate mechanical pain hypersensitivity in male and female mice. Nature Neuroscience, 18(8), 1081–1083. 10.1038/NN.4053

45. Stout, S. C., & Nemeroff, C. B. (1994). Stress and psychiatric disorders. Seminars in Neuroscience, 6(4), 271–280. 10.1006/SMNS.1994.1034

46. Teicher, M. H., Dumont, N. L., Ito, Y., Vaituzis, C., Giedd, J. N., & Andersen, S. L. (2004). Childhood neglect is associated with reduced corpus callosum area. Biological Psychiatry, 56(2), 80–85. 10.1016/J.BIOPSYCH.2004.03.016

47. Ulrich-Lai, Y. M., Figueiredo, H. F., Ostrander, M. M., Choi, D. C., Engeland, W. C., Herman, J. P., & Ulrich-Lai, Y. M. (2006). Downloaded from journals.physiology.org/journal/ajpendo at Georgia State Univ. Am J Physiol Endocrinol Metab, 291, 965–973. 10.1152/ajpendo.00070.2006.-The

48. VanRyzin, J. W., Marquardt, A. E., Argue, K. J., Vecchiarelli, H. A., Ashton, S. E., Arambula, S. E., Hill, M. N., & McCarthy, M. M. (2019). Microglial Phagocytosis of Newborn Cells Is Induced by Endocannabinoids and Sculpts Sex Differences in Juvenile Rat Social Play. Neuron, 102(2), 435–449.e6. 10.1016/j.neuron.2019.02.006

49. Watson, S., & Mackin, P. (2006). HPA axis function in mood disorders. Psychiatry, 5(5), 166–170. 10.1383/PSYT.2006.5.5.166

50. Weinhard, L., di Bartolomei, G., Bolasco, G., Machado, P., Schieber, N. L., Neniskyte, U., Exiga, M., Vadisiute, A., Raggioli, A., Schertel, A., Schwab, Y., & Gross, C. T. (2018). Microglia remodel synapses by presynaptic trogocytosis and spine head filopodia induction. Nature Communications, 9(1), 1228. 10.1038/s41467-018-03566-5

51. Yan, X.-X., Toth, Z., Schultz, L., Ribak, C. E., & Baram, T. Z. (1998). Corticotropin-Releasing Hormone (CRH)-Containing Neurons in the Immature Rat Hippocampal Formation: Light and Electron Microscopic Features and Colocalization With Glutamate Decarboxylase and Parvalbumin NIH Public Access. Hippocampus, 8(3), 231–243. 10.1002/(SICI)1098-1063(1998)8:3<231::AID-HIPO6>

52. Zhang, Y., Chen, K., Sloan, S. A., Bennett, M. L., Scholze, A. R., Sean, O., Phatnani, H. P., Guarnieri, P., Caneda, C., Ruderisch, N., Deng, S., Liddelow, S. A., Zhang, C., Daneman, R., Maniatis, T., Barres, B. A., & Wu, J. (2014). An RNA-Sequencing Transcriptome and Splicing Database of Glia, Neurons, and Vascular Cells of the Cerebral Cortex. Journal of Neuroscience, 34(36), 11929–11947. 10.1523/JNEUROSCI.1860-14.2014

